# Chronic pain alters microvascular architectural organization of somatosensory cortex

**DOI:** 10.1101/755132

**Authors:** A.G. Zippo, V. Del Grosso, A. Patera, M. P. Riccardi, I. G. Tredici, G. Bertoli, A. Mittone, M. Valente, M. Stampanoni, P. Coan, A. Bravin, G.E.M. Biella

## Abstract

Chronic pain (CP) represents a complex pathology profoundly involving both neural and glial compartments of the central nervous system. While most CP studies have also investigated the macroscopic brain vascular system, its microstructural architecture still remains largely unexplored. Further, the adaptive modifications of the vascular microstructure as consequence of diseases or pathological insults, did not receive adequate attention. Here we show microtomographic signs of diffuse and conspicuous microvascular neogenesis in somatosensory cortex of CP animal models already peaking at 15 days from the model instantiation. Progressive fading of this microvessel neogenesis then ensued in the next six months yet maintaining higher vascular density with a preserved small fraction of them. Due to the important consequences on the neuron-glial-vessel arrangements and on the resulting metabolic and functional disorders of the local networks, novel additional scenarios of CP are thus conceivable with profound consequences of potential future CP diagnostic and therapeutic appraisals.

## Introduction

At the level of the central nervous system, Chronic Pain (CP) displays a cohort of signs and symptoms, where the percept of pain dominates within a complex morpho-functional background of anomalous neuronal and glial events; these appear variously combined and causative of a global, maladaptive condition. At larger scales, brain regional anomalous activations in CP patients[2,9,59], as shown by functional neuroimaging, are seemingly accompanied by metabolic dysfunctions and local macrostructural signs such as cortical thickness reductions or cortical laminar misalignments[43]. At lower cell-level scales, CP-related functional disorders have been described, in the past, mostly as gangliar, spinal or central neuronal dynamic irregularities[7,14,51,57]; these mainly included spontaneous hyper- or hypo-activities, hyper-responsiveness to non-noxious and noxious stimuli, anomalous spiking patterns, often accompanied by variable signs of local neuronal degeneration and by cortical somatotopic remapping of sensory projection fields. These signs appeared complementary and ascribed to perceptual conditions such as spontaneous pain, allodynia and hyperalgesia.

Only recently, communal consistent complex functional network hallmarks, not simply embodied by anomalous intermittences of dynamic profiles of single neurons, have been identified in different CP models. Here, reduced neuronal information transmission and heavy topological thalamo-cortico-thalamic network rearrangements from organized to random-like graph [66] have been described. The electrophysiological dissolution of key functional traits of network connectivity it represents, however, a partial view of such a complex brain phenomenon to be analyzed on wider and more intricate background. Indeed, the tight relationships between neuronal and glial compartments and the microvascular surroundings identifies a “neuro-gliovascular unit” (NGV), a non-demountable complex with profound mutual influences in diverse functional circumstances[1,58]. Brain microvessel and capillary dynamics are now placed under deep scrutiny because of the evidences of their capital role in many brain inflammatory and degenerative pathologies[44,49,65]. Namely, where the macro- and meso-scale brain angiological connectome shows a quite stable architecture during the normal individual lifespan, microvessel or capillary niches in the brain display continuous changes of whole blood flux, blood fluid particulate dynamics, gaseous saturation due to local or external factors, activation or inactivation of subserved networks. These plastic diffuse NGV assemblage resent of major insults in pathological events, often revealed in complex patterns[45]. For instance, inflammatory episodes involving glia may be able to trigger the neo-angiogenetic processes, thus establishing a vicious cycle mutually dislodging the NGV components[61] as it has been shown in other neuropathological contexts such as Parkinson’s, Multiple Sclerosis or Alzheimer’s diseases[24,33]. Comparable studies of blood microvessel architecture and topology in CP, are, however, still lacking, missing details that could concur to elucidate and illuminate the potential role of vascular disorders on the dynamics of the central sensory networks. Here, we analyzed the somatosensory cortical microvessel architecture and topology by synchrotron micro-computed tomography (microCT) imaging in control and CP animals: our aim was to identify possible morphological and topological modifications of the somatosensory microvasculature in CP. Measures have been taken in the somatosensory cortex at three stages after the induction of neuropathy to observe fast- (2 weeks), medium- (2 months) and long- (6 months) term vessel changes, compared to same age controls.

## Methods

### Ethical Statement

All the animals (42 albino Sprague-Dawley male rats 225-250 g) have been used in accordance to the Italian, French and European Laws on animal treatment in Scientific Research (Directive 2010/63/EU on the protection of animals used for scientific purposes, transposed into the Italian legislation by the “Decreto Legislativo” of 4 March 2014, n. 26; and into the French legislation by the “Décret n° 2013-118” of 01 February 2013). The research was approved by the Italian Ministry of Health and classified as “Biella 3/2011” and further as “Biella 1/2014” into the files of the Ethical Committee of the University of Milan, Italy. For France, the authorization N. 04597.02 was provided by the French Ministry of High Education, Research and Innovation after approval by the Ethical committee #113).

### Number of Animals

We used 42 animals, randomly subdivided in three experimental groups: each group was composed by 7 Controls (CR) and 7 Constriction Injury (CP model) animals. Three groups of CP animals have been used, composed by selected at 15, 60 and 180 days from the injury date in the treated animals.

### Surgery for the neuropathic Constriction Injury model

The generation of the neuropathic constriction injury CP model was made following the (standard) protocol by Seltzer et al.[15,52]. Briefly, the animals were initially lightly anaesthetized by gaseous Isoflurane^®^ into an anesthetic chamber and then injected intraperitoneally with a solution of sodium Pentobarbital (50 mg/kg). After the full development of anesthesia, the posterior right thigh was shaved, and the skin disinfected. The rats were then placed on a heating pad, the hindpaw positioned at 90° relative to the backbone axis. A 2- to 2.5-cm long incision was performed down the center of the posterior thigh (well identifiable by finger starting from the lumbar vertebrae) parallel to the femural bone. The *biceps femoris* muscle fascicles were delicately separated by blunt-tipped surgical tweezers and the surgical wound lips of the muscle widened until a 1 cm tract of the sciatic nerve was visualized. A thin surgical forceps was used to separate in two contingents the sciatic nerve along its major axis by introducing a curved needle mounted with sterile gut suture wire into the surgical fissure within the sciatic nerve, turning the wire around the one of the two nerve halves. Without stretching the nerve, the surgical suture thread was then advanced and a tight ligature around the nerve was then performed, trimming the two wire ends at 0.5 mm above the wire knot. The muscle wound lips were then closed layer by layer with reabsorbable surgical staples. Skin suture thread was then used to close the wound. Antibiotic powder (Streptomycin sulfate salt, Sigma-Aldrich) was locally placed on the wound lips. The animals were then allowed to recover in a warm, temperature-controlled cage with a solid floor. After their complete awakening from the anesthesia, the animals were reintroduced in their original cages. In the next six hours, and up to four days after the surgery, multiple controls were done to check the normal surgical scar process and the health conditions of the animals.

### Behavioral tests

We adopted common behavioral tests[17,21] to validate the chronic pain model instances. Freely walking in a transparent plastic arena (45×65 cm^2^, with 10 cm vertical borders) allowing observation from above and from below the animals and estimations of the walking patterns, their speed and correct paw placement on the floor of the arena floor, time estimates of paw retraction-latencies during von Frey tests[17] and hot-plate[21] tests of the left posterior limb in all the experimental groups. Behavioral measurements were performed one day before, and 1, 7, 14 days after the surgical procedure, as well as the day of sacrifice.

### Perfusion method

After the loss of any reflex, before the euthanasia by the barbiturate overdose administered intraperitoneally (Pentobarbital solution, Sigma-Aldrich, 300mg/kg), the animals were intracardiacally perfused according to a common protocol[4]. The procedure was performed in four stages: heparinized saline (150-180 ml), 10% formalin (150 ml), air (ten minutes of air insufflation through the same route of fluids and, eventually, undiluted Indian-ink (100 ml), this last used in accordance with the modified protocol by Xue et al.[64]. The unusual step of injecting air into the circulating system after the vessel fixation was adopted to empty the vascular system before the ink injection. The perfusion of the solution of heparinized saline was started after waiting few seconds for the right atrial enlargement (due to the circulatory volume expansion). The ink perfusion (Pure Rotring^®^ Ink, Rotring, Germany) was simply obtained by placing the tubes into the ink solution. A clear sign of the complete perfusion was the generalized blackening of all the mucosae, of the nude skin surfaces (such as the snout, the paws), the thoracic viscera and the eyes.

The animal sacrifice was performed at three different time stages (15 days, 2 months, and 6 months) after the sciatic ligature and the creation of the CP neuropathic model.

### Cortical sample preparation

At the established time stages (15 days, 2 months, and 6 months), the perfused animals underwent to brain extraction in order to collect samples of somatosensory cortex by coring the cortical surface by means of a cylindrical Teflon tube (2.1 mm in diameter and 12 mm in height). The tube was positioned with one end positioned in correspondence to the somatosensory region and delicately but firmly pushed into the brain until reaching the surface of the brain base, by gently turning it through the brain tissue, until the basis of the brain was reached. The obtained sample was immersed in polyphosphate buffered solution (PBS, ThermoFisher Scientific, United Kingdom) and maintained in constant hydration conditions until the microCT analysis. After the samples were used for microCT, they were embedded in paraffin blocks.

Paraffin blocks were sectioned by a microtome (LEICA RM2125) in order to obtain 5 µm thick slices. After cutting, the sections have been placed on the surface of the 37 °C water bath and then floated onto the surface of clean glass slides. The operation was repeated until all the tissue slices were gathered. The slides were dried overnight at 37 °C and stored at room temperature.

### Immunofluorescence Imaging

We applied immunofluorescence imaging to qualitatively assess the presence of vascular neo-genesis in somatosensory cortical samples. It consisted in several consecutive steps here illustrated.

#### Tissue inclusion in Paraffin

According to standard protocols, brain tissue was dehydrated through a series of graded ethanol baths to displace the water inside the tissue. Indeed, the tissue, after being washed in PBS, was maintained 18 hours in ethanol (EtOH) 70% in a gentle agitation (Reciprocating Shaker Labline, LabX, Canada, 15-20 motions per minute). Then, changing the bath each hour, it was passed through EtOH 80%, 95% and two times in EtOH 100%. After the dehydration procedure, the tissue was put in a clearing agent (Xylene) for 1 hour to displace the ethanol, and then 1 hour in Xylene-paraffin (1:1) at 56°C. Then, the tissue was embedded in a Tissue Mold (Leica Biosystems, USA), covered with paraffin in a liquid state and let it becoming cold. When the wax was completely cooled and hardened (30 minutes), the paraffin block was popped out of the Tissue Mold.

#### Deparaffinization

The paraffin was removed from the slides by rinsing them for 5 minutes in Xylene, 10 minutes in EtOH 100%, 5 minutes in EtOH 95%, 5 minutes in EtOH 75%. Then, they were rehydrated by immersion in deionized water at first, and then in PBS, five minutes each.

#### Antigen retrieval

According to the datasheet of the used primary antibody (CD34, Abcam, Cambridge, United Kingdom), formalin-fixed tissue requires the antigen retrieval step before immunohistochemical staining. During this step the disulphide bridges were broken and so the protein antigenic sites were easily exposed to antibodies binding. A solution of sodium citrate (10mM, pH 6.0) was prepared and heated up to 98 °C. The slides were immersed here for 3 minutes and then they were left cooling down for 40 minutes at room temperature. Finally, they were washed 3 times in PBS for 2 minutes each under a gentle agitation (15-20 motions per minute).

#### Permeabilization and Saturation

The antibody entrance in the cell was promoted in this process by producing microlesions on the cell surfaces with Triton^®^ X-100 solution (Sigma-Aldrich, USA). A solution of 0.2% Tween20-PBS was used. The slides were maintained in this solution for 30 minutes under a gentle agitation (15-20 motions per minute). Saturation was needed to block unspecific binding of the antibodies, thus, a solution of 10% bovine serum albumin (BSA, Sigma-Aldrich, catalog A2058) in 0.1% Triton-PBS was used. The slides were maintained for 1-2 hours at 4 °C under gentle agitation (15-20 motions per minute).

#### Immunofluorescence staining

The slides have been incubated in a solution containing the primary CD34 antibody (Abcam, Cambridge, United Kingdom, catalog ab185732) (1:500 in BSA 10%) for 1 hour at room temperature. Thereafter, the solution was decanted and the slides were washed in PBS three times, for 5 minutes each, in gentle agitation. The slices were then immersed in a solution containing goat anti-mouse secondary antibody (GAR 488, Abcam, Cambridge, United Kingdom, catalog ab150077) (1:2000) for 1 hour at room temperature in a dark environment.

Then, the slices were incubated with Hoechst33342 (Abcam, Cambridge, United Kingdom, catalog ab145597) in PBS (1:500) for 5 minutes, still in a dark environment, and rinsed with PBS for 5 minutes. Hoechst stains are part of a family of blue fluorescent dye used to stain the DNA of cell nuclei. Coverslips were mounted on the slides with a drop (5 μl) of mounting medium (30% glycerol in PBS) and sealed with nail polish to prevent drying and movement under microscope. A Nikon Eclipse80i microscope equipped with mercury vapor lamp has been used. The used filters were DAPI/Hoechst filter (340-380nm excitation, 435-485nm emission) and FluoresceinIsothiocyanate (FITC) filter (465-495nm excitation, 515-555nm emission). The pictures were taken with automatic exposure by the Nikon software called ‘NIS elements’ (Nikon OS.02/L2 USB; F package).

### Scanning Electron Microscopy (SEM)

Textural and morphological observations (secondary electron images) were performed by means of a Tescan FE-SEM (Mira 3XMU-series, Tescan, Czech Republic), equipped with an EDAX energy-dispersive spectrometer for compositional microanalysis. SEM imaging was performed with an In-Beam high resolution detector, at different magnifications, ranging from 100× to 20k×. The samples were the same tissue slices on glass slides described in the previous section.

Before observation, the samples were coated with Pt using a (Cressington, United Kingdom) 208 HR sputter (deposition time: 60 s, deposition current: 20 mA).

An accelerating voltage of 8 kV and 20 kV was applied for morphological observation and composition microanalysis, respectively. The latter was performed with counts of 100 s per analysis. The measurements were processed using the EDAX Genesis software (Tescan, Czech Republic).

### Statistical Analyses

For group comparisons we considered the Kruskal-Wallis non-parametric statistical test[34] while multiple pairwise differences were calculated by the Tukey’s post-hoc Honest Significant Difference test[60]. Singular pairwise comparisons were done by the Wilcoxon non-parametric rank sum test[63]. The significance level was set to 0.05 in all analyses.

### Synchrotron radiation propagation-based phase-contrast micro-Computed Tomography imaging

The microCT images were acquired at the TOMCAT beamline of the Swiss Light Source, Paul Scherrer Institute, Villigen, Switzerland[19,48]. The X-ray source is a 2.9 T superbending magnet with a critical energy of 11.1 keV[19]. The quasi monochromatic (few percent bandwidth) X-ray beam (energy: 20.0 keV) was extracted by Ru/C multilayer mirror. The sample was placed on a remotely controlled rotation stage placed at 20 mm from the source, which is an optimized distance for recording propagation-based phase contrast images. X-rays passing through the sample reach a LSO:Tb scintillator of 5.9 µm in thickness converting X-rays into visible light, before being collected by a diffraction-limited microscope optics with a selected 20× objective (NA=0.70); the image is then projected on a pco.edge sCMOS camera with a 16-bit dynamic range and a field-of-view (FoV) of 832×702 µm^2^ on the horizontal and vertical directions, respectively; the effective final pixel size is 0.325 µm.

For each tomography, 1401 (plus 30 dark fields and 100 flat fields) radiographic projections were acquired at equiangular positions over a total rotation angle range of 180° with an exposure time of 300 ms each. Four stacked scans were acquired along the vertical direction of the sample in order to image at least 2 mm over 8 mm of cortical cortex sample, resulting in a total scanning time of 30 minutes per sample.

Acquired images presented a marked edge enhancement at the borders of the details, which is the effect of the propagation-based phase-contrast imaging setup used for image acquisition combined with the X-ray beam with high degree of spatial coherence available at TOMCAT[12].

The radiographic projections were processed using the Paganin algorithm[47], before being reconstructed using the *gridrec*[41] algorithm. All 3D renderings were made using the ImageJ software.

### Vessel Segmentation

While cardiac perfusion with Indian-ink efficiently marked blood vessels[37,64], the extraction of their putative regions within the tomographic volumes remained a non-trivial operation. A Python toolbox was developed ad-hoc to gather blood vessel voxels by a sequence of three incremental steps: i) an opportunely tuned Canny edge detector algorithm[13] extracted the initial set of voxels; ii) morphological operators (dilation, filling and erosion) refined the initial set; iii) an unsupervised clustering method drastically reduced false positive and negative items. Details of the implementation choices and source code are available at https://github.com/antoniozip/xray_vessel_segmentation.

### Complex Brain Network

For the topological analyses, the vascular system has been *skeletonized* (a modeling technique which transforms vessels in collections of cylinders eventually collapsing into *diameterless* rods) and then characterized by the number of skeleton segments, the number of branch points, the vascular length and the maximum flow in order to investigate potential changes in CP condition. Topological analyses converted the entire vascular structures extracted from tomographies into large single graphs with tens of thousands of nodes, one for each experiment. Vascular networks have been analyzed with a couple of common complex network statistical metrics, namely the clustering coefficient and the characteristic path length[62]. The former returns an estimation of the network ability to process information in segregated clusters of node, the latter measures the network ability to spread information among nodes. A further characterization from a graph-theoretical perspective has been performed by estimating the maximum flow among network nodes. Estimations of the maximum flow between all network node couples were performed by an open source Python implementation of the Edmonds-Karp algorithm[18] (https://github.com/bigbighd604/Python/tree/master/graph).

## RESULTS

We used an experimental and computational framework to reconstruct the 3D microvascular system of cortical brain samples (**Figure 1**) from control (CR) and neuropathic (CP) animal groups, the latter analyzed at three different time phases of the neuropathy: 2 weeks, 2 months and 6 months (see Methods for further details). Qualitatively, the analysis of the 3D images returned a clear scenario where CP samples were much more vascularized than control (**Figure 2**). This was particularly evident in the 2-week model and progressively reducing in the 2-month and 6-month samples.

**Figure 1.**
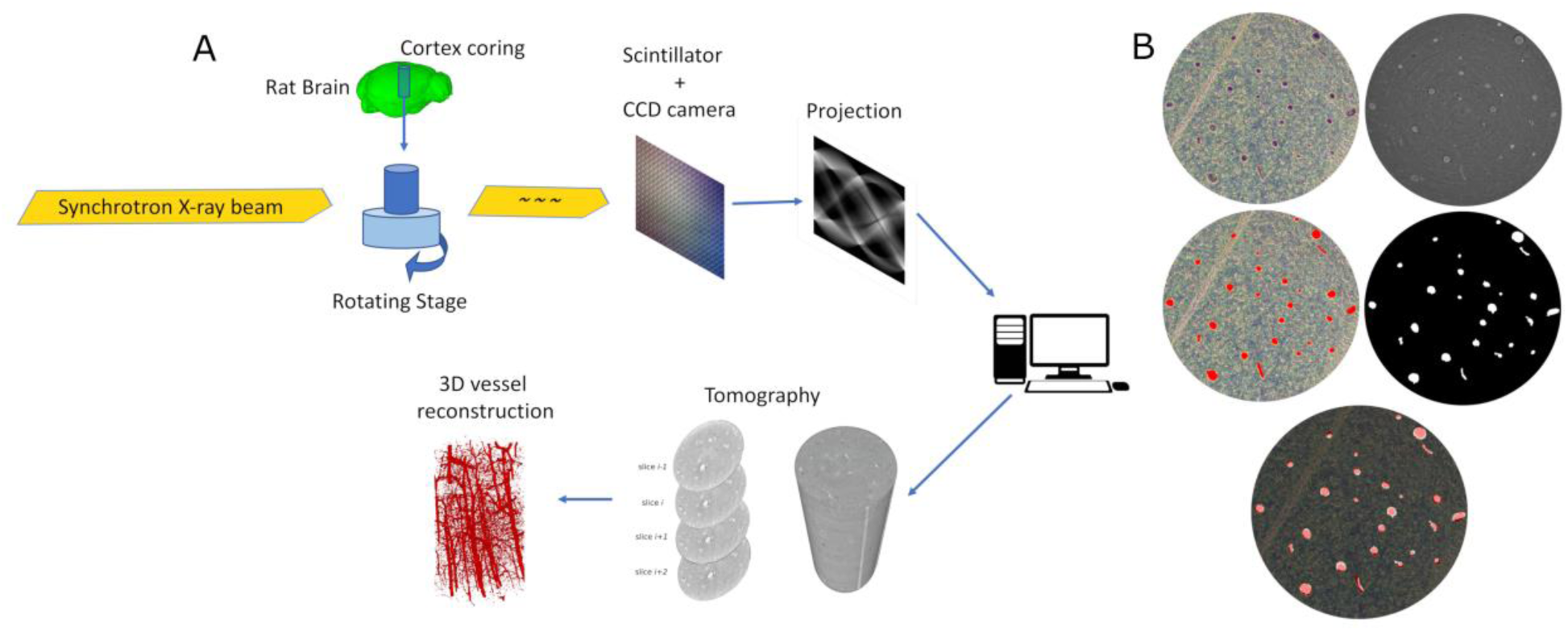
The experimental framework used to reconstruct the 3D vessel microstructure from the cortical coring (A). The synchrotron-generated X-ray beam hit the cortical sample held on a rotating stage. The resulting beam excites a scintillator followed by a digital camera sensor. Resulting projections are processed by a computer that reconstructs the 3D microCTs slice-by-slice. At last, an automatic segmentation algorithm extracts the entire 3D vascularization of the scanned brain tissue sample. (B) Histological validation of X-ray microtomographic microvessel extraction. A sample of histological slice where microvessels are highlighted by Indian-ink (black, first column, first row). In the right column of the same row, is displayed the equivalent slice obtained by X-ray microCT. In the second row (left), manual segmentation of microvessels in the histological sample. Same row, on the right: automatic segmentation of microvessels in the microtomographic sample. The last slice on the bottom shows the merged slice that match between the histological and putative algorithmically extracted microvessels.

**Figure 2.**
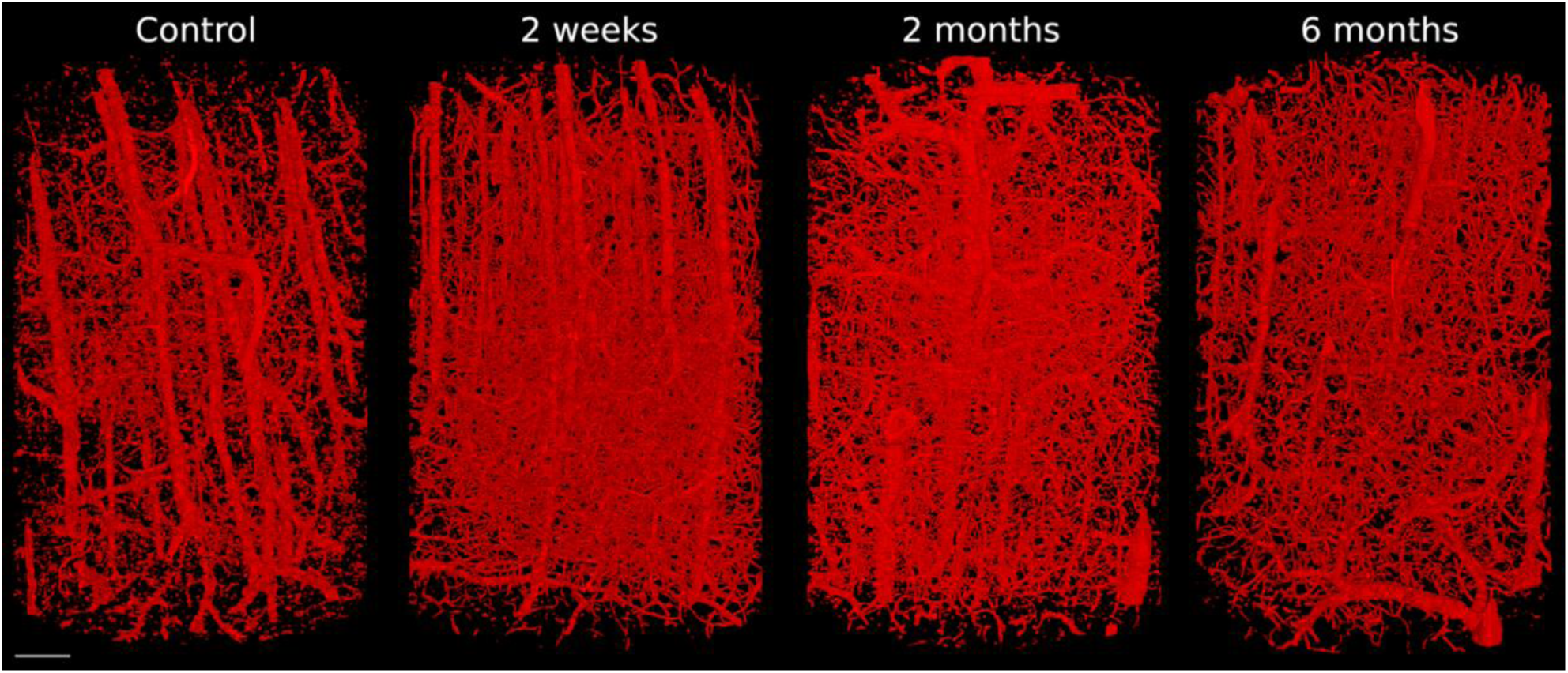
Comparison among 3D reconstructed vascular systems sampled from the four experimental conditions (Control, 2 weeks CP, 2 months CP and 6 months CP). Chronic pain models appeared much more densely vascularized. Scale bar is indicated on the bottom white line that corresponds to 100 *μ*m.

### Morphological comparisons

Morphometrical investigations took into consideration two crucial features of the vascular system: vessel diameter and density. In the former, the average vascular diameter was estimated through geometrical assumptions for each sample; in the latter we computed the percentage of volume occupied by vessels in the total volume. We found that vessel diameters were significantly smaller in the 2-week and 2-month models than in controls and in 6-month samples (Kruskal-Wallis test with Tukey post-hoc Honest Significant Difference test, significances in **Figure 3A**). Similarly, the vessel density was, in general, much higher in CP models than in control animals with a marked emphasis in the 2-week and 2-month subgroups, which partially vanished in the 6-month one (**Figure 3B**). Differences in density were also factorized for cortical layers (I, II/III, IV and Va). We found that layers contributed non-homogeneously in the vascular density distribution (**Figure 3C-D**) in all CP groups. Furthermore, Tukey post-hoc Honest Significant Difference test (**Figure S1**) revealed that Layer I represented the main target for the angiogenesis and their maintenance, a phenomenon progressively less evident in deeper layers. In detail, while the 2-week group showed no specificity of layer, the 2-month models, instead, showed a higher density for layers from I to IV and not for layer Va. In the 6-month models, the vascular density increment was only visible in layer I while layers IV and Va had significantly lower vascular densities.

**Figure 3.**
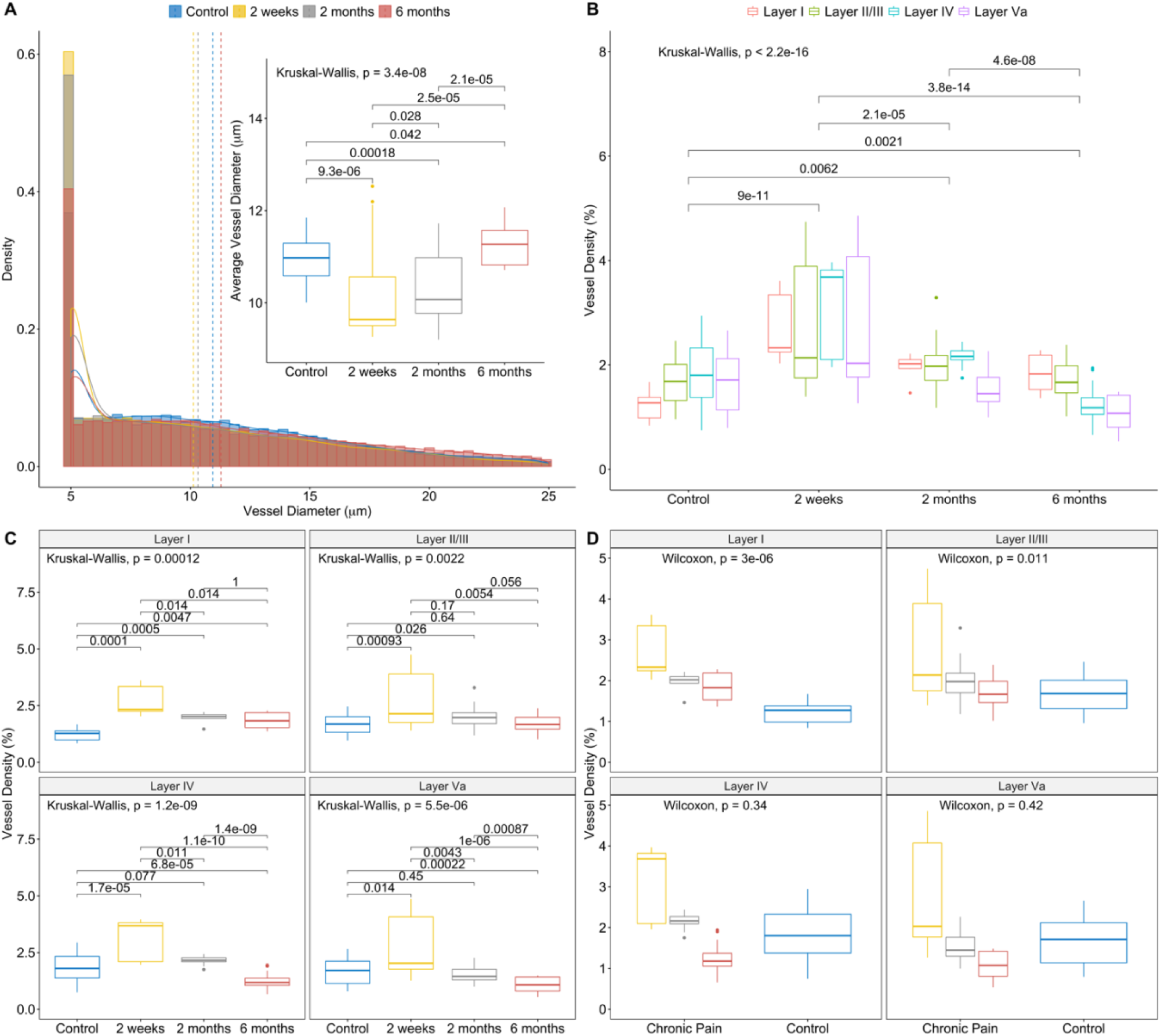
Estimation of the vessel diameter and density distributions in the four experimental groups and cortical layers. (A) The dashed lines represent the estimated expected values. The continuous lines approximate the probability density distribution for each group. Histograms are cut off at 25 *μ*m because no differences are visually appreciable. The inset shows the same distributions with box-whisker plot layout where the internal horizontal three lines represent respectively the 25^th^, 50^th^ and 75^th^ percentiles and the line extremes the 1^st^ and 99^th^ percentiles. Pairwise significances were obtained by Tukey post-hoc test for differences. (B) Box-Whisker plots of the distribution of the vessel diameters in control and the three CP conditions divided in the four cortical layers. (C) The four experimental conditions were analyzed layer-by-layer. (D) The 2 main group (control and chronic pain) were compared layer-by-layer.

Statistical hypothesis tests were consequently affected by a small population size effect (N = 10) and although distributions were visually much different, they rarely reached the significance level (0.05). We found that CP induced an increment of the number of branch points in comparison to controls (**Figure 4A**, Kruskal-Wallis tests and Tukey post-hoc tests for pairwise differences). Specifically, taking into account the three different time stages, the increments appeared consistent (although not significant with the Wilcoxon non-parametric rank-sum test) at two weeks, two months and six months. This result indicated the alteration of the branch point structure during the stabilization of CP in the somatosensory cortical tissue. Since branch points are the originating points of bifurcations, the CP onset modified the vascular network producing novel vessels originating from the extant structure. Besides the findings on branch points, comparable dynamics have been observed also for the number of segments in the vascular skeletonized version. Indeed, the number of segments grew up in CP (**Figure 4B**). The **Figure 4C** visualizes the estimated length of the skeleton segments along the four experimental conditions. Again, we found that in control animals, vessels were much shorter than in the 3 CP conditions (**Figure 4C**).

**Figure 4.**
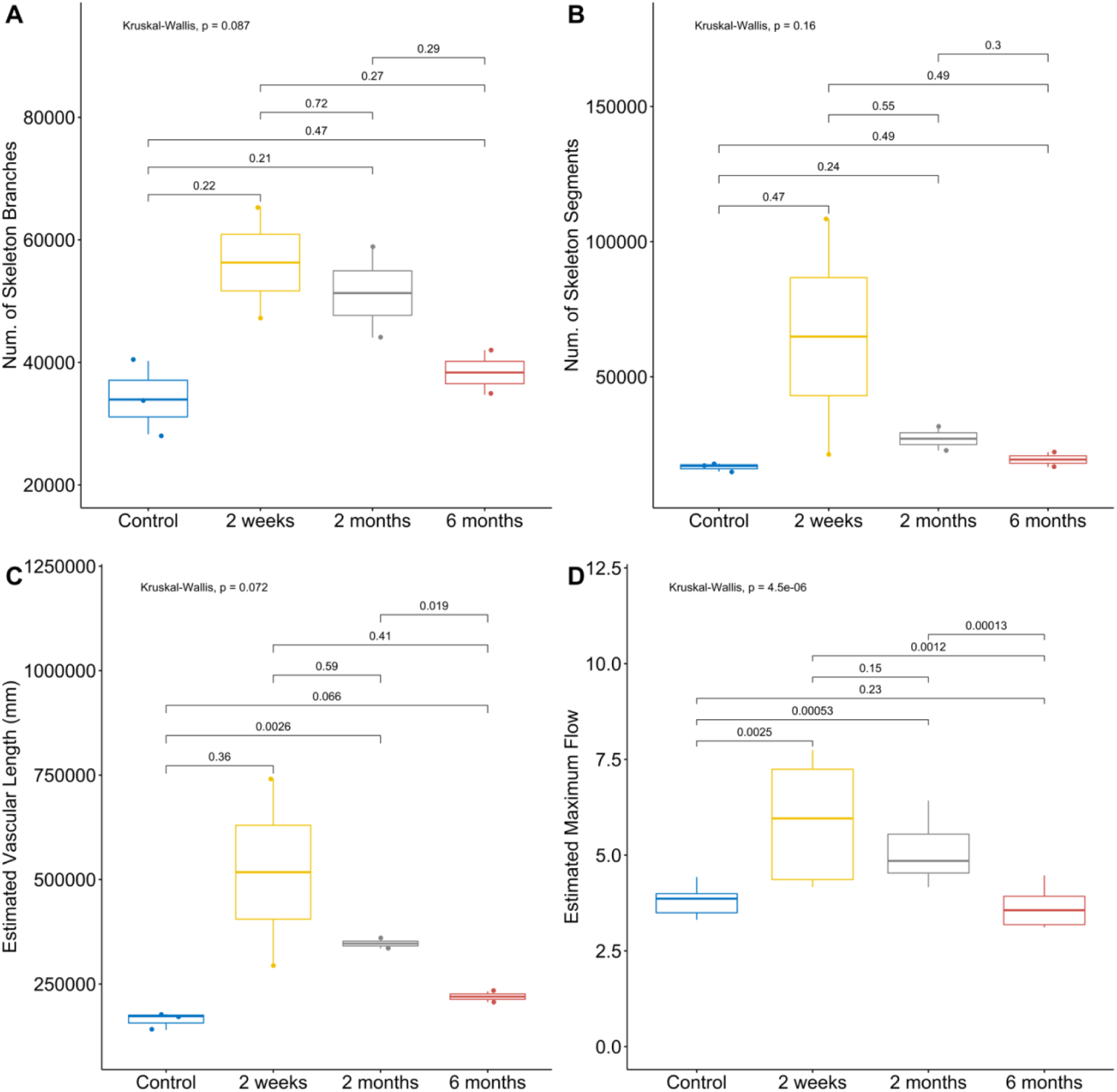
Vessel topological features computed on the skeletonized version of the microvascular network. (A) Number of skeleton branches (A) and segments (B). Estimated vascular lengths in mm (C) distribution of maximum graph flow on the skeleton (D). All distributions are represented by Box-Whisker plots (1^st^ percentile, 2^nd^ quartile, median, 3^rd^ quartile, 99^th^ percentile; dots represent outliers) measures were computed in the four experimental conditions control (blue), 2 weeks (yellow), 2 months (gray) and 6 months (red). Pairwise significances were obtained by Tukey post-hoc test for differences.

### Topological characterizations

Summarily, the requirement of a greater number of segments to describe the CP vascular network suggested that CP vascular networks had a much larger and more complex morphology. Additionally, to extend the theoretical interpretation of observed results, a graph theory metrics has been used. Namely, the *maximum flow* evaluated the theoretical capacity of a network (modeled as a *pipeline*) to load low viscosity fluids (e.g. water or blood). We found that the maximum flow sustainable in CP vascular networks was much greater than control networks (**Figure 4D**). Specifically, 2 weeks after the CP model inset, the vascular networks reached the maximum network flow, followed by the maximum flow of the 2-month rat models; controls, instead, had higher maximum flow than 6-month rat models. Since maximum flow estimation did not consider the speed of flow, assumed under constant pressure, this result indicated that CP vascular networks were compatible with an enriched blood flow sustained by the promoted novel angiogenesis.

### Immunostaining and SEM analyses

Eventually, we performed a last set of experimental analyses to investigate the molecular substrate of the observed neoangiogenesis. We chose as target protein the transmembrane phosphoglycoprotein CD34, normally expressed by blood vessel endothelial cells. Immunofluorescence imaging showed a significant higher concentration of microvessels in CP models than in control (**Figure 5**, a result also appreciable in bright field in **Figure S2**), with a vanishing dynamic comparable to the analytical results: a peaking effect at 2 weeks, progressively fading at 2 months, and then at 6 months. As an additional experimental control, we performed SEM imaging of the histological slices of the cortical coring acquired by synchrotron microCT,. This control assessed that the contrast-medium Indian-ink effectively permeated vessels and that the resulting imaging was in accordance with our previous results. **Figure 6A** shows a comparison among the 4 experimental conditions with SEM images acquired with a small magnification. The different concentration of microvessels (radiopaque white spots) can be seen also at higher magnification (**Figure 6B**). Finally, a spectrographic analysis of an Indian-ink filled microvessels revealed its carbon-rich molecular fingerprint (**Figure 6D**, different from the rest of the brain tissue (**Figure 6E**).

**Figure 5.**
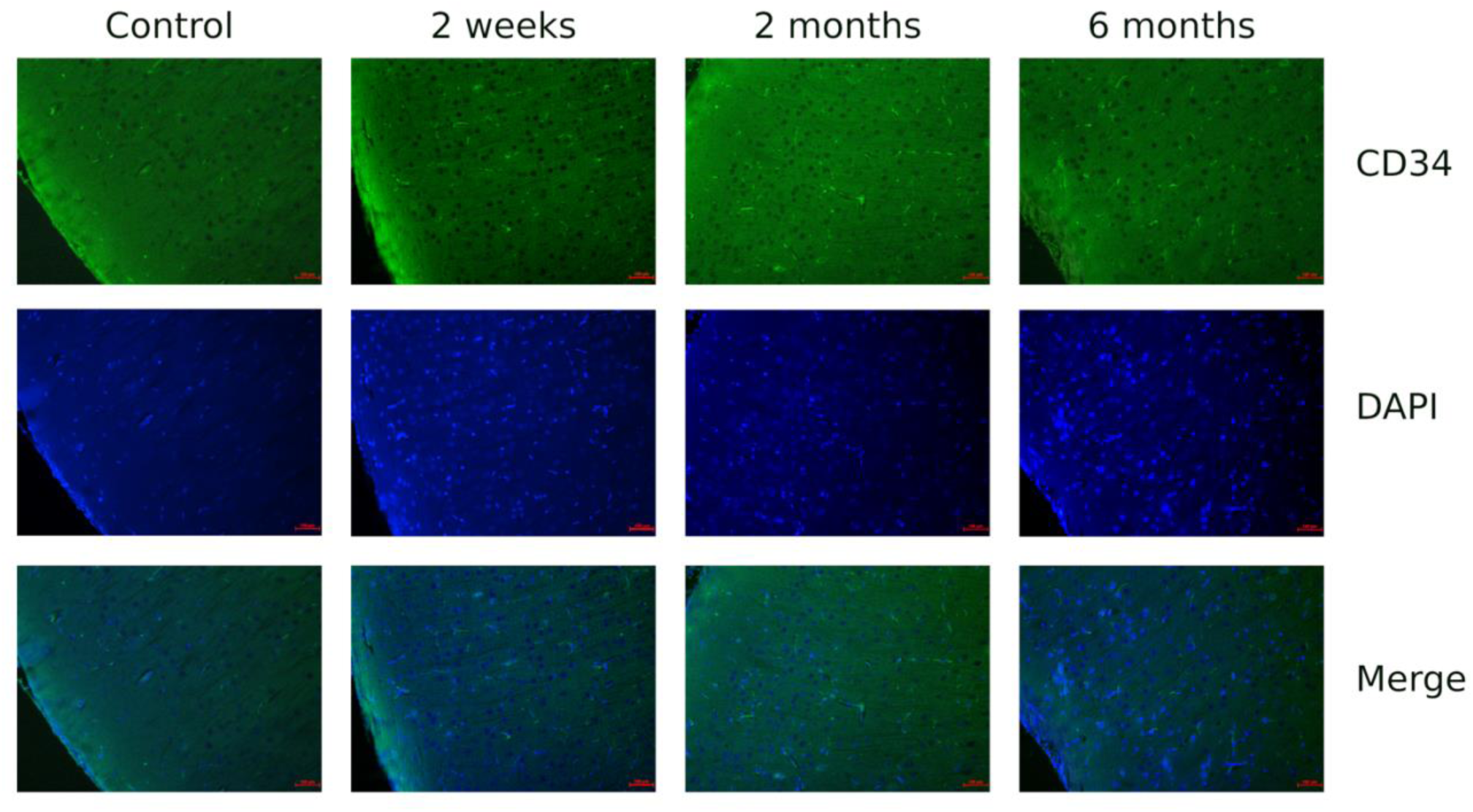
Immunofluorescence analysis of coronal slices sampled from each group. Tissues in the first row are treated primarily with an anti-CD34 antibody targeting the vascular endothelial cells, combined with the Alexa Fluor^®^ 488 secondary antibody. The second row slices were stained with Hoechst 33258 staining dye solution targeting the cell nuclei. The third row displays the corresponding merge between the first and the second row images. Scale bar are plotted in red at the bottom right border of each slice (100 *μ*m).

**Figure 6.**
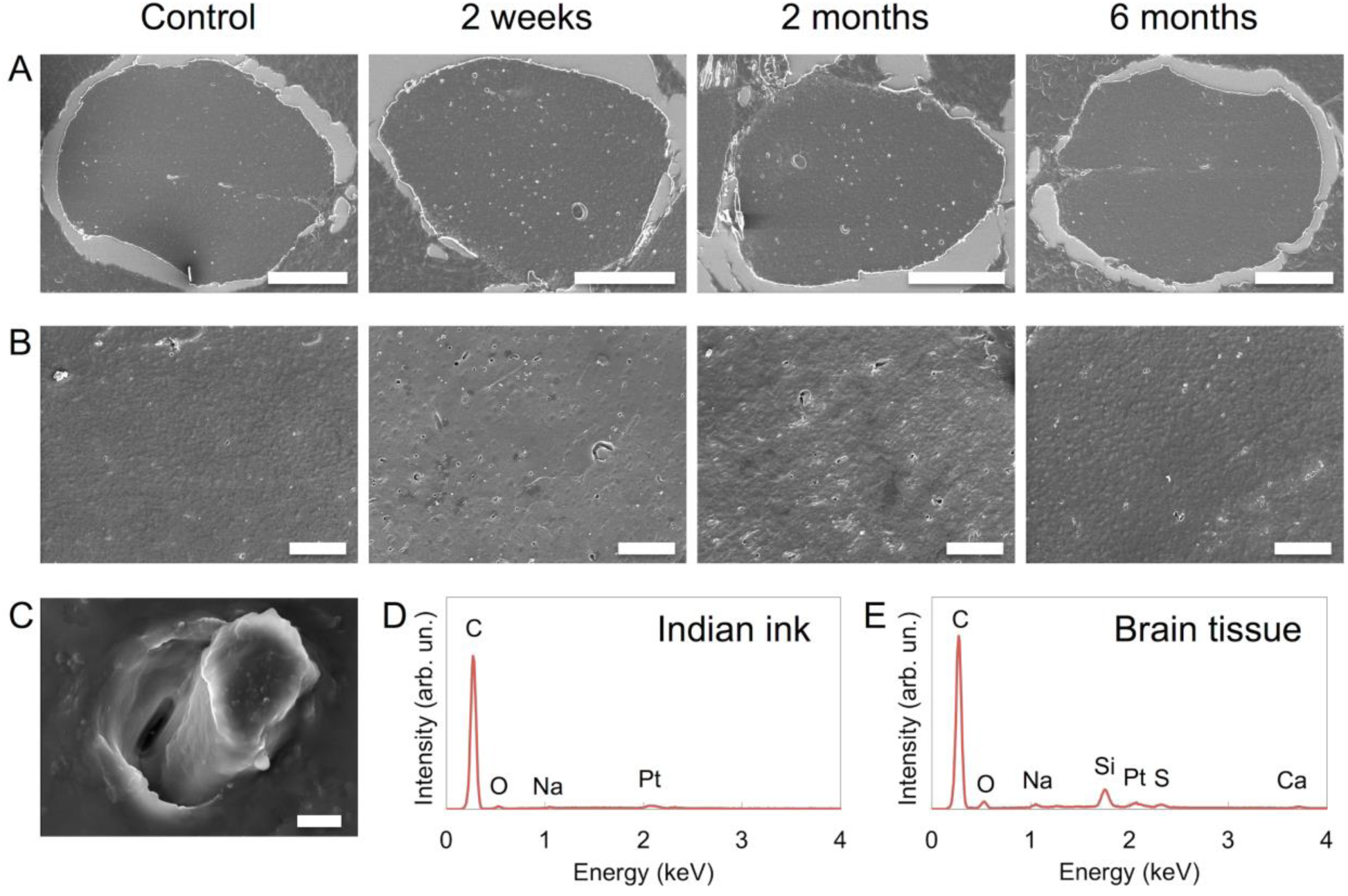
Scanning Electron Microscopy Imaging. (A) Large field images of the histological slices (6 *μ*m thick) obtained by cortical samples used in the X-ray synchrotron microCT; vessel walls are highlighted in white. Scale bars: 500 μm. In (B) images were registered at higher magnification (500x, scale bars 100 μm). (C) A large magnification (20 kx) of the solidified Indian-ink within a microvessels of ∼ 5 *μm* in diameter (scale bar: 2 μm). A spectral confirmation of the presence of Indian ink is represented by the spectrogram in (D) that emphasizes the copious and almost exclusive presence of carbon, in comparison to a spectrogram extracted from a non-specific point of the tissue (E) where other biological elements are present.

Altogether, these results suggested that the primary somatosensory cortex is subject to a conspicuous short-term neoangiogenesis of microvessels, as a result of a peripheral neuropathy. Furthermore, topologically abnormal microvessels were produced, and although compensatory dynamics tried to reestablish the proper architectural regime, long-lasting fallouts were stabilized after 6 months from the initial neuropathy injury.

## DISCUSSION

In this paper we show, by microtomographic imaging acquired at a synchrotron radiation X-ray beamline, a widespread numeral increase of blood microvessel and capillaries in the somatosensory cortical region of interest of an experimental rat model of chronic pain. The microvessel proliferation was particularly remarkable just in the early stages of the pathology (at 2 weeks) showing in the next six months a progressive reduction of this upsurge yet maintaining traces of hyper-vascularization (at six months, Figure 2, fourth column). This phenomenon has been interpreted as a consequence of the neuronal metabolic needs from anomalous activities in CP where changes in microvessels may indeed not only respond to but also sustain the alterations of the served neurons.

The brain microCT imaging has already been applied in comparable experimental contexts where quantification methods at the mesoscale on brain cortex have provided isotropic resolution images of the tissue, thus revealing 3D architectures not only of vessels but also of neuroglial compartments[22]. Comparisons with classical SEM have also shown not only a perfect tissue preservation after microCT sessions but also a straightforward correspondence in co-registration analyses between the two techniques. This assessment ensures the reliability of this new technique as standard procedure in brain microstructural studies[6,20].

It is valuable here to consider all actors: the pre- and post-synaptic neuronal terminals, the astrocytic synaptic envelopes and the related microvessels which build up a hardly dividable *functional entity* or neuro-glial-vascular (NGV)-*unit*. Each element of this unit is deeply involved in the global dynamics of the unit itself. As a matter of fact, all the disorders involving even only one element of the *NGV unit* appear to implicate all the other unit elements[30,31]. Our results could help to examine, in a new perspective, the hemodynamic conditions expressed in CP and conjecture, along with the progressive pathogenesis, the ensuing metabolic costs and the subsequent complex adaptive responses of the NGV unit in CP genesis and maintenance[35]. As it was found by Jensen[28],, in normal conditions, the vessel compartments match the neuronal density variety throughout the cortical laminae[28] and that no particular branching or vessel density variation occurs in time. Beyond the structural data, other issues rise about the functional efficiency of these neogenic vessels as well as about their comparability with the extant microvessel assembly. Although our computational estimation of the *maximum flow*, it remains to be clarified if an efficient blood flux in the anomalously packed microstructures may take place, yet globally preserving stable biophysical processes such as blood shear thinning or thixotropy, i.e. the blood fluidity changes with shape changes, or the gaseous and solute exchanges. They should obey also to Murray’s law[46] that states how, in vascular systems, daughter vessel diameters are related to parental vessels in a regime where the radii of the vessels must be in relation to flow (the larger the radius, the smaller *power* required for *flow*, Pf), and to metabolic upkeep costs or *power* of the vessel *wall tissue* (Pwt). Namely, vessels have to be neither too large nor too small if the *total power*, Pt = Pf + Pwt, is to be minimized, a configuration probably acquired under selective demands[53]. Remarkable discrepancies, however, have been noticed among the neogenetic events in the different cortical layers. Unexpectedly, the first layer showed a longer lasting permanence of increased vascularization in comparison with other inspected layers: it could be due to the higher number of local synapses originating from lower layers and in particular to the extensive dendritic trees of lower layers [16]. The slow but significant, even if not complete, decay of the microvessel density surge in the following months may, then, be interpreted as an adapting phenomenon, or a recovery process of exceeding upkeep costs of the newly generated vessels and the sustenance of pathological CP-related energetic costs[8,25,32,54–56].

In topological terms, the unchanged values of clustering coefficient and the characteristic path length suggest that the surge of novel microvessels, as well as the decay of the majority of them in the subsequent months, present a steady topological organization. As a side note, the vascular event recoverable in CP patients by functional Magnetic Resonance Imaging gross inspection, as mosaic activation-inactivation, may become more complex to be understood, being assignable also to microvessel volume changes with more debatable signs of activations of already extant vessel compartment[3,5,42].

From a functional point of view, a large array of mechanisms of transcytotic and transcellular transport of gases and various substances and nutrients characterizes the rich interactions among glial perivascular endfeet and microvessels [27]. The neo angiogenetic microvessels, to be effective, should indeed establish mutual efficient contacts with the glial interface. From our observations and estimates on increased whole blood fluxes, it is conceivable that at least a significant fraction of these vessels may represent active energy sites for different reasons. In first place, the surge of the phenomenon could be hardly denied in a period when high neuronal energy consumption, with additional energetic costs of vessel neogenesis, would have no energy trade-off counterbalance[40]. Secondarily, the novel vessels could play protective roles to avoid that severe energy shortage conditions might trigger hypoxic and apoptotic processes[36]. As final observation, all these neogenic events could not the chaotic outcome of irregular events dictated by an extreme condition such as CP but emerge as modifications with the same topology of the control networks. This is remarkable in that kind of local stable *Bauplan* that seems inherent to the vessel arborization and independent from the pathological originating conditions. The slightly supernumerary small vessel contingent remaining at six months could be probably considered a long-term remnant response of metabolic compensation, integrated within the brain tissue on a hyperactive or anomalous neuronal network. A further question may be raised on the percentage of open vessels in the diverse brain compartments. This complex phenomenon depending on the hypoxic, hypercapnic, energetic factors, as with adenosine dynamic, as well from intracranial pressure and blood brain barrier permeability, may be left in the background in the experimental conditions we operated in. In fact, all the animals have been perfused in following steps with pressures higher than the common rat vascular blood pressure measured around 70 mm Hg in normal conditions[11]. In our system, indeed, the output pressure through the cardiac intraventricular (left ventricle) needle was about 150 mmHg with a higher velocity flow in comparison to natural parameters. As it has been shown[10], there are two classes of high and low velocity, where proportion of low-velocity capillary (<1 mm/sec, 3–7 µm diameter) appears to represent the 67.2%, significantly greater than the high-velocity microvessels, with a percentage of 32.8% (>1 mm/sec, 8–15 µm diameter). However, the pump perfusion pressures were seemingly able to cancel all the fine dynamics of the brain microvessels, thus imposing a global high pressure regime disrupting the fine details of the varieties of the diverse hemodynamic regimes, as shown in works on pathological experimental pressures, where wide recruitment of vessel opening was recorded[11,26]. Capillaries are a group of small diameter vessels that are grossly distinguished by their diameters from shunts by their size (3–7 μm diameter), branching, and tortuosity. Larger-diameter microvessels from 8 μm to 45 μm are in the range of arterioles and venules and arterio-venous, arterio-arterio, and veno-veno shunts that are precapillary shunts capable of shunting blood away from capillary beds. Though this variety of elements in the microcirculatory context, due also to the absolute replication of the perfusion methodology in the different rat models and in the diverse time evolution of CP models, we are confident that, also from a theoretical point of view, none of these reported elements could weaken the assumption of the numeral increase and subsequent decrease of the capillary bed elements in our experimental conditions.

Our method presents also some advantage in tomographic 3D resolution in comparison with the recently presented automated protocol for measuring the capillary diameter[38] and also with the brain capillary reconstruction with the CLARITY method that in planar reconstruction achieves a pixel resolution of 0.65 μm, needing however 2 mm thick slices[23].

At last, very recent findings obtained by anti-Stokes Raman scattering microscopy have shown that Aquaporin-1 provides mechanical static or pulsed-pressure induced deformation protection of vessels[39,50]. Disruption of this mechanism appears contributing to endothelial dysfunction in various disease states and to inflammatory local responses, in turn sources of vessel neogenesis. Chronic pain often accompanies inflammatory episodes in the brain[29] and these inflammatory seeds may represent a further push toward a positive milieu for the microvessel and capillary growth with the cohort of related anomalies. It can be hypothesized, that the prominent and fast neoangiogenesis in altered sensory conditions might promote immature structures of vessels that, together with metabolic factor, favor the degeneracy of newborn vessels. In conclusion, a novel actor of the nervous tissue appears dramatically involved in the chronical instantiation of CP in the somatosensory cortex. envisaging various interventions of the potential role of angiogenesis inhibitor agents for the treatment or prevention of CP in human subjects.

## Acknowledgments

We wish to thank Ms. Chiara Bellei and Dr. Fabio Zucca from the Institute of Biomedical Technologies (Consiglio Nazionale delle Ricerche) and Dr. Alexandra Demory for their support and assistance in histological sectioning and paraffin inclusion of brain samples; the team of the ID17 beamline and of the Biomedical Facility (ESRF) for sample preparations and data processing; the TOMCAT personnel (PSI) for the support provided in the realization of beamtimes. Authors acknowledge support provided by the SYRA3 COST Action TD1205 (AB, VDG, GEMB, PC), by the BIONECA COST Action CA16122 (AB) and by the Deutsche Forschungsgemeinschaft (Cluster of Excellence) - Munich Center for Advanced Photonics EXE158 (PC, AB). The authors have not conflict of interest.

## Author contributions

G.B. conceived the experiment. A.Z., A.P., A.B., M.S. V.G. and P.C. actively led experimental sessions. A.Z. performed data analyses, statistical tests and produced all images. M.R. and I.T. performed electron microscopy imaging. A.Z. and GB performed immunostaining experiments. A.Z., A.B. and G.B. wrote the manuscript. All authors contributed to the manuscript revision.

## List of Figures

**Figure S1.**
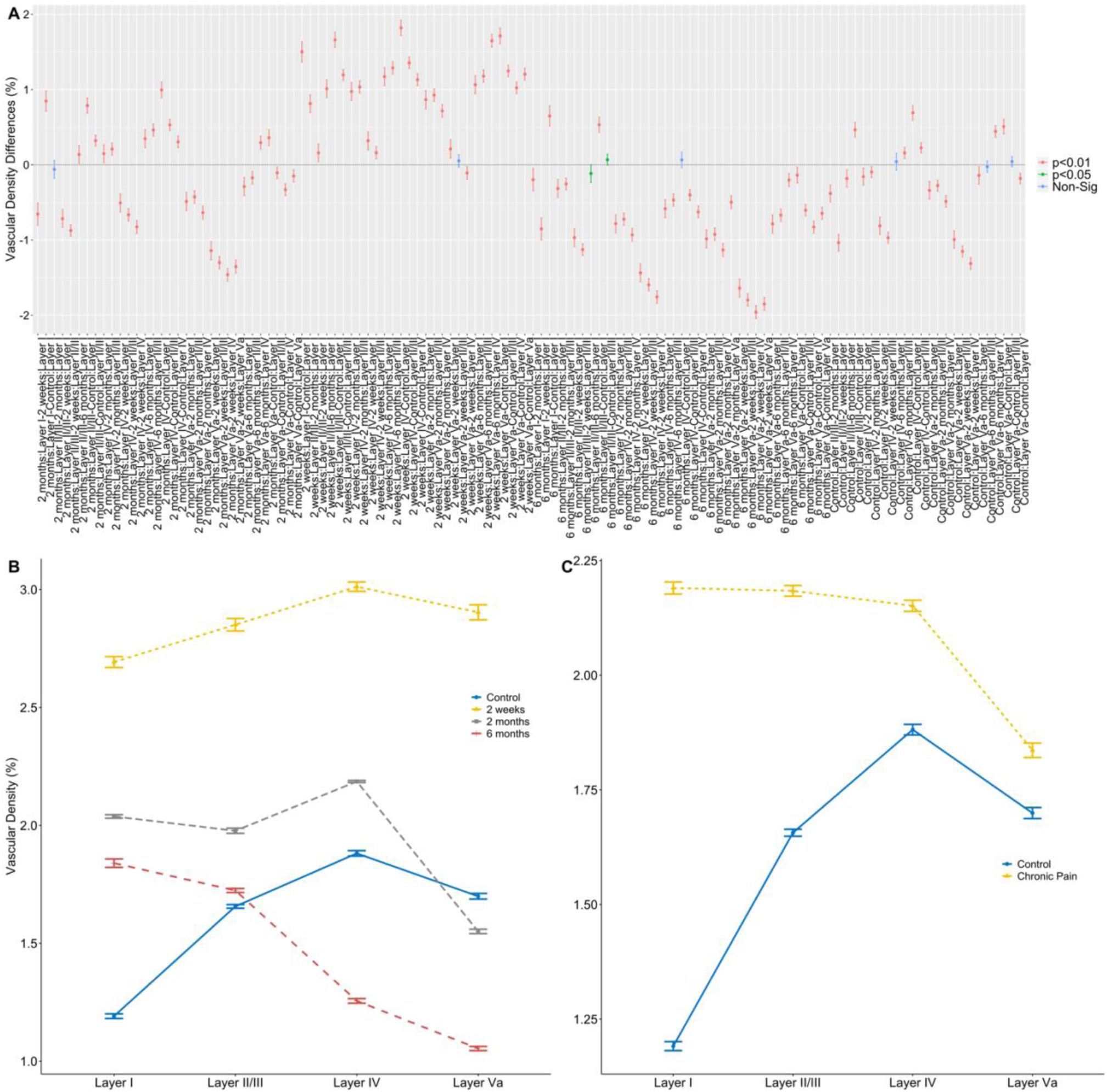
Additional analyses of the distribution of vascular density among groups and layers. (A) A representation of the post-hoc analysis (Tukey’s test with a confidence interval of 0.95) which contrasts each layer with each group. (B) Interaction plot of the vascular density between cortical layers and groups. (C) Interaction plot of the vascular density between cortical layers and the major experimental conditions (control, chronic pain). In (A) filled circles indicate the mean value and the bars (A, B, C) the standard deviations.

**Figure S2.**
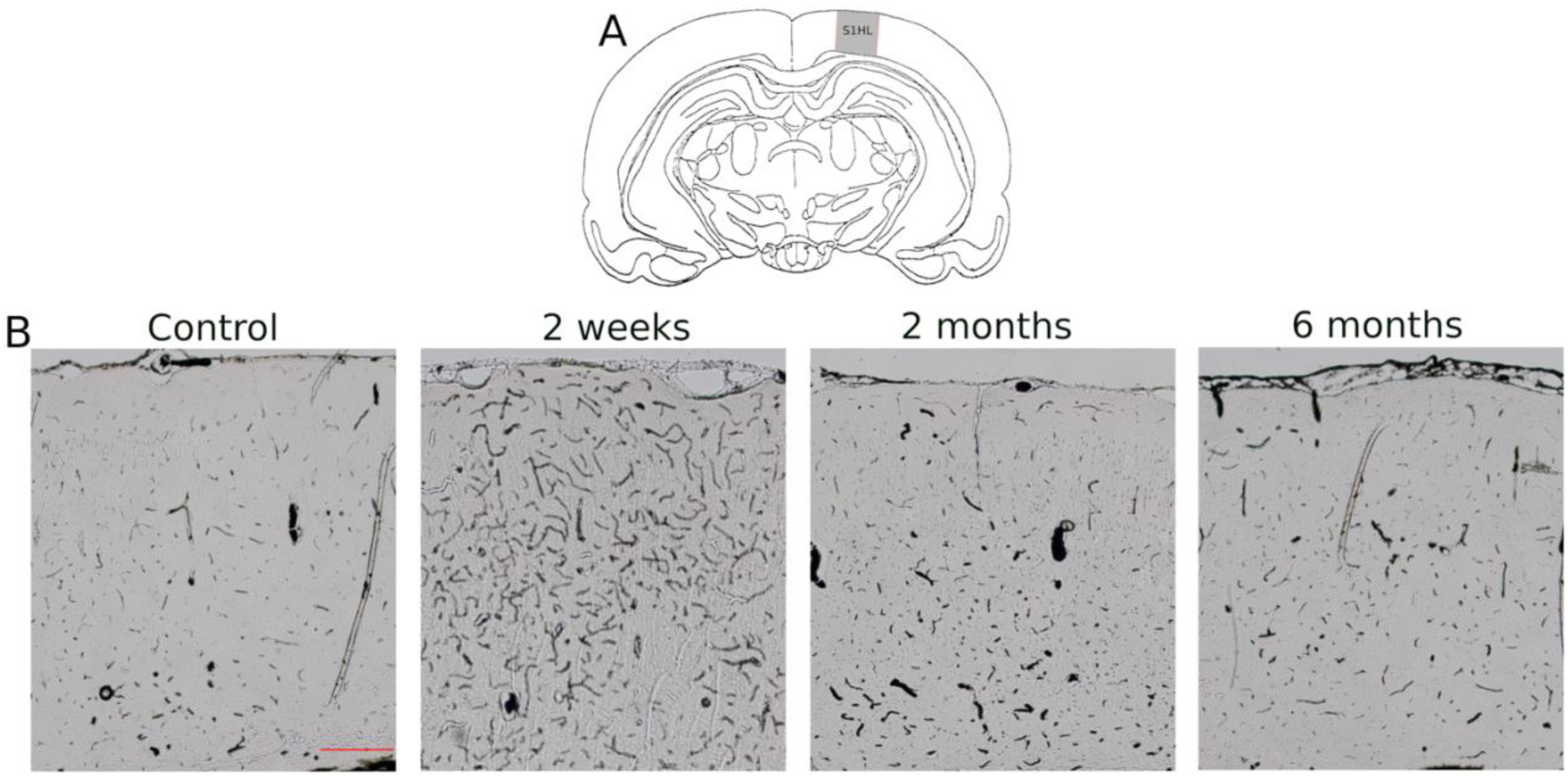
Bright field acquisition of histological samples from the 4 experimental group. (A) The schematic coronal section of the regions of interest, i.e. the primary somatosensory cortex (S1), hindlimb projection (HL). (B) A qualitative comparison of the deparaffinized slices where black thread indicated the Indian-Ink perfused vessels. Scale bar in the control column is 400 µm.

## List of Movies

**Movies S1-4**. 360-degree rotation of the reconstructed microvascular cortical cylinders in the four groups: CR (S1), CP at 2 weeks (S2), CP at 2 months (S3) and CP at 6 months (S4).

